# AGO1 interacts with *NEAT1* lncRNA and impacts nuclear compartments

**DOI:** 10.1101/2025.01.26.634929

**Authors:** Muhammad Shuaib, Taro Mannen, Tomohiro Yamazaki, Sabir Abdu Adroub, Yanal Ghosheh, Waad Albawardi, Tetsuro Hirose, Valerio Orlando

**Affiliations:** King Abdullah University of Science and Technology (KAUST), BESE Division, KAUST Environmental Epigenetics Program, Thuwal, 23955-6900, Saudi Arabia; College of Life Sciences, Ritsumeikan University, Kusatsu 525-8577, Japan; Graduate School of Frontier Biosciences, Osaka University 1-3 Yamadaoka, Suita, Osaka 565-0871, Japan; IRCSS Fondazione Santa Lucia, Epigenetics and Genome Reprogramming, Rome, Italy

**Keywords:** Argonaute-1 (AGO1), *NEAT1* lncRNA, Paraspeckles, Chromatin Compartments

## Abstract

Argonaute 1 (AGO1), a central component of the RNA interference (RNAi) pathway, has recently been implicated in nuclear functions, particularly in chromatin organization and gene regulation. However, the role of AGO1 in lncRNA-driven nuclear compartmentalization remains elusive. In this study, we uncover a novel function for AGO1 in the regulation of nuclear architecture through its interaction with the long non-coding RNA (lncRNA) *NEAT1*. *NEAT1* lncRNA is required for the formation of paraspeckles, nuclear substructures involved in multiple cellular processes. We show that AGO1 physically interacts with *NEAT1* and other key paraspeckle proteins (PSPs) and co-localizes with paraspeckles in the nucleus. AGO1 depletion disrupts expression of both *NEAT1* isoforms, reduces the interaction of essential PSPs with *NEAT1* lncRNA, and leads to impaired paraspeckle formation. Furthermore, depletion of *NEAT1* results in the mis-localization of AGO1 from paraspeckles and alters active chromatin compartments. Together, our findings establish a new functional relationship between AGO1 and *NEAT1* in nuclear compartmentalization and chromatin architecture.

## Introduction

Argonaute proteins (AGO1-4) are key players in the RNA interference (RNAi) pathway, mediating post-transcriptional gene silencing primarily in the cytoplasm through their interactions with small RNAs (Joshua-Tor and Hannon, 2011; Meister, 2013). Among these, Argonaute 1 (AGO1) has recently been identified as having nuclear functions (Allo et al., 2014; Huang et al., 2013) that extend beyond its canonical RNAi activity. Our previous work demonstrated that nuclear AGO1 binds to active chromatin, playing a critical role in chromatin organization and gene expression regulation in human cells (Shuaib et al., 2019). However, the molecular details of how AGO1 operates within the nuclear landscape, particularly its interactions with long noncoding RNAs (lncRNAs) and nuclear substructures, remain poorly understood.

Among the several lncRNAs discovered, Nuclear Enriched Abundant Transcript 1 (*NEAT1*) has emerged as an important player in nuclear organization. *NEAT1* is required for the formation of paraspeckles, which are subnuclear structures involved in multiple processes within cells such as gene control, stress response, and development (Rinn and Chang, 2012). The *NEAT1* gene produces two transcripts: *NEAT1_1*, a 3.7-kb transcript, and *NEAT1_2*, a longer 23-kb transcript, among which *NEAT1_2* but not *NEAT1_1* is essential for paraspeckle assembly (Clemson et al., 2009; Mao et al., 2011). Paraspeckle formation is initiated by the transcription of *NEAT1*, followed by its interaction with over 50 paraspeckle-associated proteins (PSPs), which include RNA-binding proteins and other nuclear factors (Chujo et al., 2017; Naganuma et al., 2012; Sasaki et al., 2009). Although the precise biological function of paraspeckles remains under investigation, growing evidence indicates that they regulate gene expression by sequestering RNA and proteins within the nucleus (Hirose et al., 2014; Imamura et al., 2014; Prasanth et al., 2005). The *NEAT1* lncRNA has also been shown to associate with active genomic regions and promote phase separation-mediated paraspeckle assembly (West et al., 2014; Yamazaki et al., 2018). This evidence points to a broader role for *NEAT1* as an architectural RNA to maintain intranuclear architecture and gene expression, although its potential involvement in organizing chromatin domains has yet to be fully explored.

In this study, we identify nuclear AGO1 as a novel paraspeckle component that controls *NEAT1* expression and affects paraspeckle assembly. We show that AGO1 directly interacts with both *NEAT1* isoforms and essential paraspeckle proteins (PSPs) and co-localizes with paraspeckles within the nucleus. AGO1 depletion not only disrupts *NEAT1* expression but also diminishes its interaction with main PSPs, resulting in defective paraspeckle formation. Notably, the loss of *NEAT1*, along with the resulting disappearance of paraspeckles, disrupts AGO1 localization and impacts chromatin organization, especially concerning A/B compartmentalization. This supports the notion that AGO1 and *NEAT1* lncRNA work together to regulate nuclear architecture (Shuaib et al., 2019). Collectively, our findings provide new insights into the functional interplay between AGO1 and *NEAT1* architectural lncRNA in shaping subnuclear structure in human cells.

## Results

### Chromatin-bound AGO1 RIP-seq identifies *NEAT1* lncRNA as a major partner

Our previous studies have shown that RNA facilitates the connection of AGO1 with active chromatin and its role in gene regulation (Shuaib et al., 2019); nevertheless, its direct interaction and functional relationship with nuclear-enriched lncRNAs remains unexplored. To identify enriched ncRNAs, we examined chromatin-bound AGO1 RIP-seq data. The RIP-seq was performed in two RNA size fractions (100 nt and 400 nt), both of which displayed similar profiles (Figure 1A). The genome-wide mapping of the AGO1-associated RNA fractions (∼100 and ∼400 bp) showed enrichment of coding, non-coding and un-annotated RNA peaks (Figure 1B-C). We identified a total of 3500 and 1854 transcripts containing approximately 50% non-coding RNAs in both 100 bp and 400 bp RIP-seq fractions respectively.

**Figure 1.**
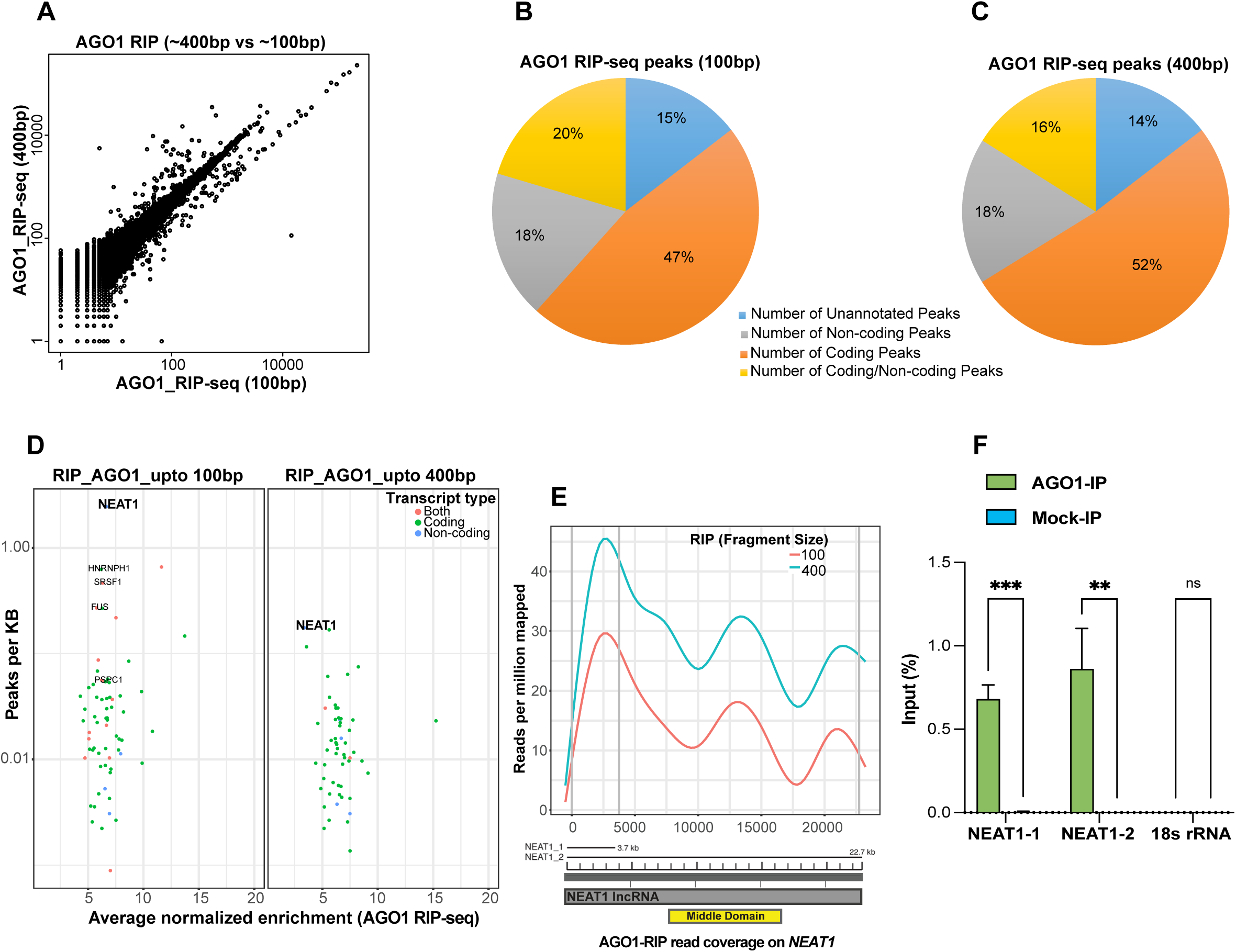
Analysis of chromatin-bound AGO1-associated RNAs. (**A**) AGO1 RIP-seq from chromatin-bound RNA fractions (∼100 bp and ∼400 bp). Mapping sequencing reads from these two fractions display similar distributions. (**B-C**) Pie-charts show AGO1 RIP-seq peaks classification identified after mapping and peak-calling using Piranha (V1.2.1) in the two fractions: (**B**) ∼100 bp (**C**) ∼400 bp. (**D**) Representation of highly enriched AGO1 associated chromatin bound transcripts (coding, non-coding and both). (**E**) Read coverage enrichment on *NEAT1* lncRNA (short and long isoforms) and its essential middle domain in AGO1 RIP-seq fraction (100 bp and 400 bp). (**F**) Validation of direct interaction between AGO1 and *NEAT1* lncRNA by UV cross-linking IP. After UV cross-linking, the RNAs immunoprecipitated with AGO1 IP were quantified by qRT-PCR, and the RNA enrichment (as % input) was determined (± SD from 3 experiments). IgG IP was used as mock control.

Among AGO1-associated lncRNAs, we found *NEAT1* lncRNA as the most significantly enriched major partner of AGO1 in both RNA fractions (Figure 1D). We also detected transcripts of other paraspeckle-associated proteins, including PSPC1, FUS, and HNRNPH1 (Figure 1D). AGO1 RIP-seq showed enrichment across both *NEAT1* isoforms, with a notable binding on the middle domain of *NEAT1_2* isoform (Figure 1E), which is necessary and sufficient for de novo assembly of paraspeckles(Yamazaki et al., 2018). The direct interaction between AGO1 and *NEAT1* isoforms (*NEAT1_1* and *NEAT1_2*) was further validated by UV cross-linked immunoprecipitation followed by qPCR (Figure 1F). These results suggest a functional link between nuclear AGO1 and *NEAT1* lncRNA containing paraspeckles.

### AGO1 as a novel paraspeckle component that interacts with essential paraspeckle proteins

Next, we sought to verify whether AGO1 is an integral component of paraspeckle nuclear bodies. We performed RNA immuno-FISH to visualize the localization of *NEAT1* lncRNA and AGO1 within paraspeckles. Immunofluorescence combined with RNA FISH revealed AGO1 co-localization with *NEAT1* lncRNA containing paraspeckles across three cell types: HeLa, HepG2, and HAP1 (Figure 2A-C). The quantification of RNA immuno-FISH data demonstrated that AGO1 is present in more than 50% of paraspeckles in each of these cell types (Figure 2D), indicating a strong association between AGO1 and *NEAT1*-containing paraspeckles. It is important to note that AGO1 is also detected in some other nuclear foci that are distinct from paraspeckles.

**Figure 2.**
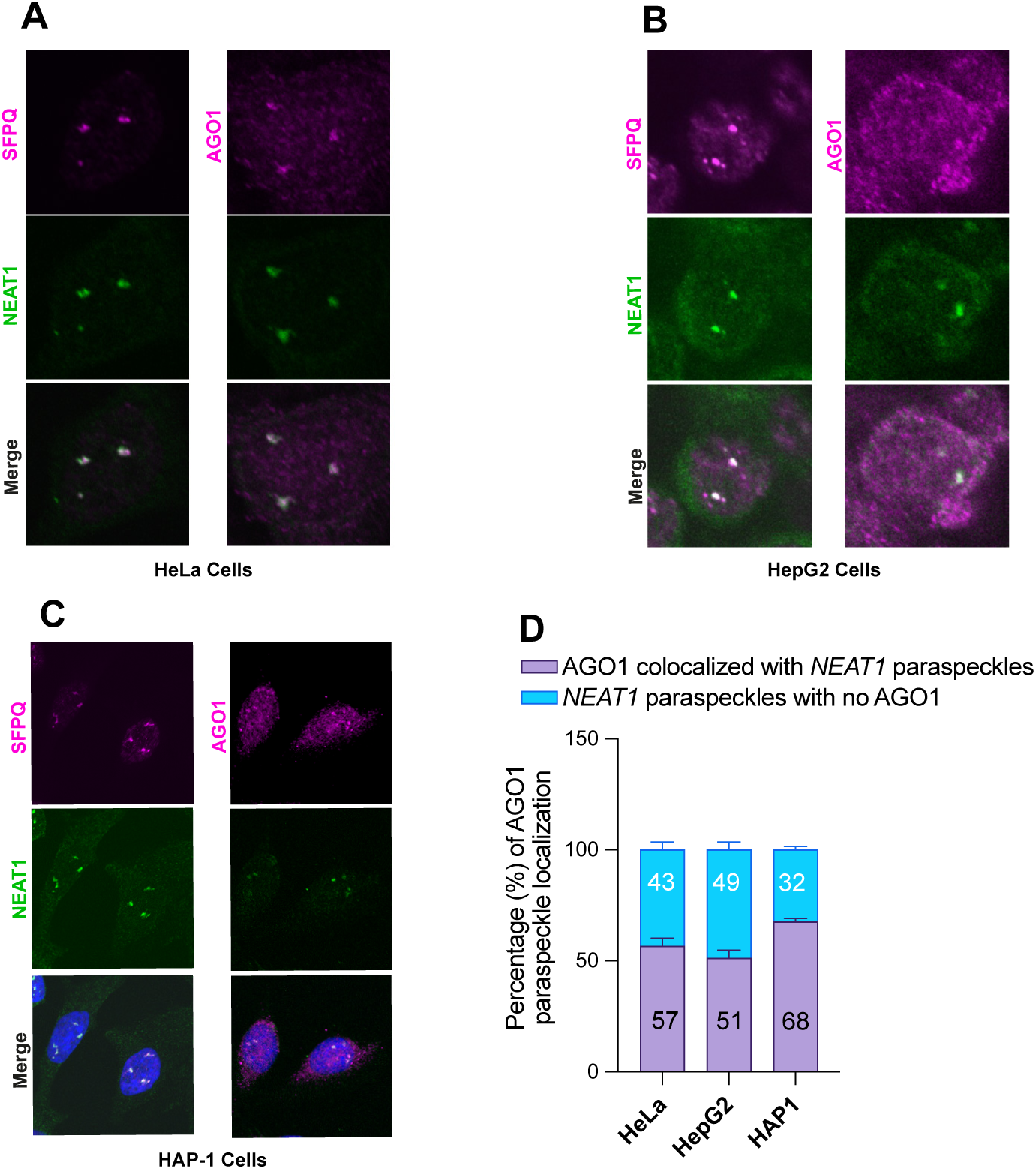
Paraspeckle localization of nuclear AGO1. (**A-C**) Localization of AGO1 and SFPQ, a known interactor, relative to paraspeckles in human (**A**) HeLa, (**B**) HepG2 and (**C**) HAP1 cells. AGO1 and SFPQ were detected by immunofluorescence and *NEAT1* RNA-FISH was used to visualize paraspeckles. (**D**) Quantification of AGO1 localized paraspeckle (n=100) in HeLa, HepG2 and HAP1 cells.

Given the observed localization of AGO1 with *NEAT1*-containing paraspeckles, we wanted to examine whether AGO1 also associates with essential paraspeckle proteins. To this end, we used nuclear and chromatin extracts of stable cell lines expressing AGO1 and AGO2 fused with C-terminal Flag- and HA-epitope tags (named e-AGO1 and e-AGO2) for double-immunoaffinity purification. Proteins associated with e-AGO1 and e-AGO2 nuclear soluble (S) and chromatin bound (CB) complexes were separated by SDS-PAGE gel (4–12%) and visualized by silver staining and western-blot (Figure 3A-B). Several proteins were found to be physically interacting with e-AGO1 and e-AGO2 soluble and chromatin complexes (Figure 3A-B). Mass spectrometry analysis allowed the identification of many paraspeckle proteins including the key components (SFPQ, FUS, RBM14, and HNRNPK) explicitly in the e-AGO1 but not e-AGO2 nuclear and chromatin complexes (Figure 3A-C). Western blot of the purified complexes confirmed the specific association of essential paraspeckle proteins (SFPQ, NONO, FUS, and HNRNPK) with e-AGO1 (Figure 3D), suggesting that AGO1 function as a new component of paraspeckle.

**Figure 3.**
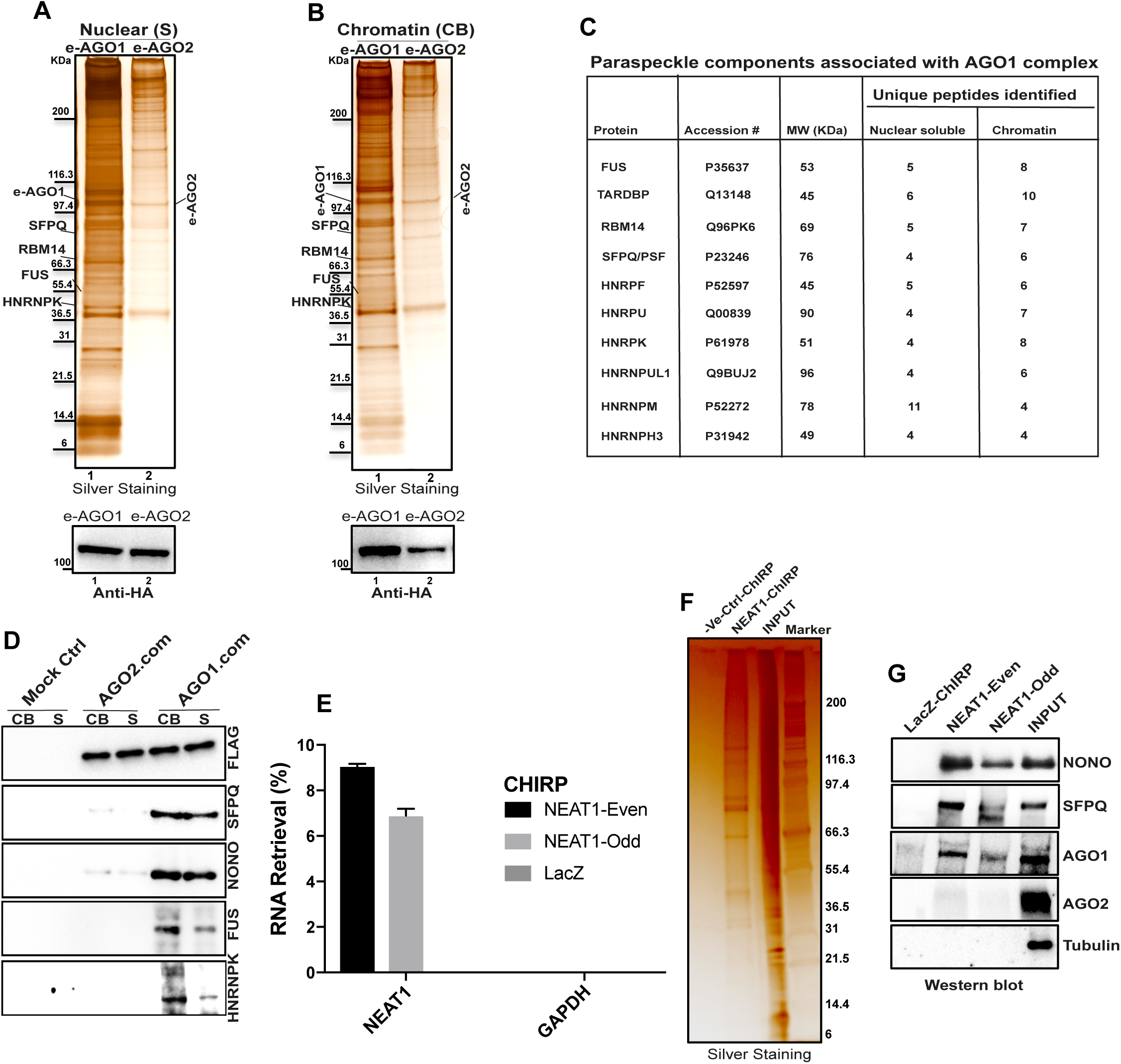
AGO1 interacts with essential paraspeckle proteins. HEK-293 cell lines stably expressing FLAG-HA epitope tagged e-AGO1 and e-AGO2 were established. After fractionation into soluble nuclear and chromatin the e-AGO1 and e-AGO2 protein complexes were purified by double affinity purification, separated on SDS-PAGE and then subjected to silver staining, mass spectrometry analysis, and western blotting. (**A-B**) Silver staining (top) of e-AGO1 (lane1) and e-AGO2 (lane2) associated proteins from (**A**) nuclear soluble and (**B**) chromatin fractions. Protein bands were identified by mass spectrometry analysis and the positions of molecular size markers are indicated. Paraspeckle components were identified in e-AGO1 nuclear and chromatin complexes. Western blot (lower) using HA antibody shows level of e-AGO1 (lane1) and e-AGO2 (lane2) in (**A**) nuclear soluble and (**B**) chromatin complexes. (**C**) Paraspeckle proteins identified by mass spectrometry analysis in e-AGO1 complexes together with the number of identified peptides are indicated. (**D**) The e-AGO1, e-AGO2 and mock control (Mock ctrl) nuclear soluble (S) and chromatin (CB) complexes (AGO1-com and AGO2-com) were analyzed by immunoblotting with the indicated antibodies. Essential paraspeckle proteins SFPQ, NONO, FUS, and HNRNPK are specific to e-AGO1 nuclear and chromatin complex. Double immunoaffinity was performed on untagged HEK-293 nuclear and chromatin extract as a mock control (Mock ctrl). (**E**) *NEAT1* ChIRP-Western blot was performed using *NEAT1* probes (odd and even) and negative control LacZ probes. Isolated RNAs were analyzed by qRT-PCR using specific primers for *NEAT1* and GAPDH (negative target). Bar chart shows successful *NEAT1* lncRNA retrieval (%) with both odd and even *NEAT1* probes (± SD from 3 experiments). (**F**) Proteins retrieved by *NEAT1* lncRNA and negative control probe along with input protein are visualized by silver staining. (**G**) Immunoblotting of *NEAT1* ChIRP-enriched proteins (isolated with odd and even pool) using specific antibodies to detect NONO, SFPQ, AGO1, AGO2, and Tubulin.

We next conducted *NEAT1* ChIRP-western blot experiment to confirm the interaction of AGO1 with *NEAT1* (Figure 3E). Many proteins associated with *NEAT1* lncRNA were separated by SDS-PAGE and visualized by silver staining (Figure 3F). Immunoblotting of the *NEAT1* ChIRP associated proteins further proved the presence of AGO1 along with essential paraspeckle proteins (SFPQ and NONO) (Figure 3G). These results indicate that AGO1 is stably associated with paraspeckle.

### *NEAT1* depletion impacts AGO1 localization

To understand the functional link between *NEAT1* and AGO1, we asked whether *NEAT1* is necessary and sufficient for AGO1 nuclear paraspeckle localization. First, we examined the nuclear localization of AGO1 upon transient depletion of *NEAT1* by antisense oligo (ASO-*NEAT1*) that targets both *NEAT1* isoforms. Upon *NEAT1* depletion, AGO1 signals, like SFPQ disappeared compared to control (ASO-GFP) treated cells (Figure 4A-C). Next, we used *NEAT1* knocked-out (KO) human HAP1 (near-haploid) cell line, generated by CRISPR/Cas9 (Yamazaki and Hirose, 2021). By combining *NEAT1* RNA FISH with SFPQ immunofluorescence, we confirmed the complete disappearance of paraspeckles in the selected *NEAT1*-KO HAP1 cell lines (Figure 4D). Consistent with *NEAT1* transient depletion, we observed a substantial adverse effect on AGO1 nuclear localization in *NEAT1*-KO cells (Figure 4E-F). *NEAT1* depletion is known to completely block paraspeckle assembly, and the absence of AGO1 signals under these conditions further confirms AGO1 association with *NEAT1*-containing paraspeckles.

**Figure 4.**
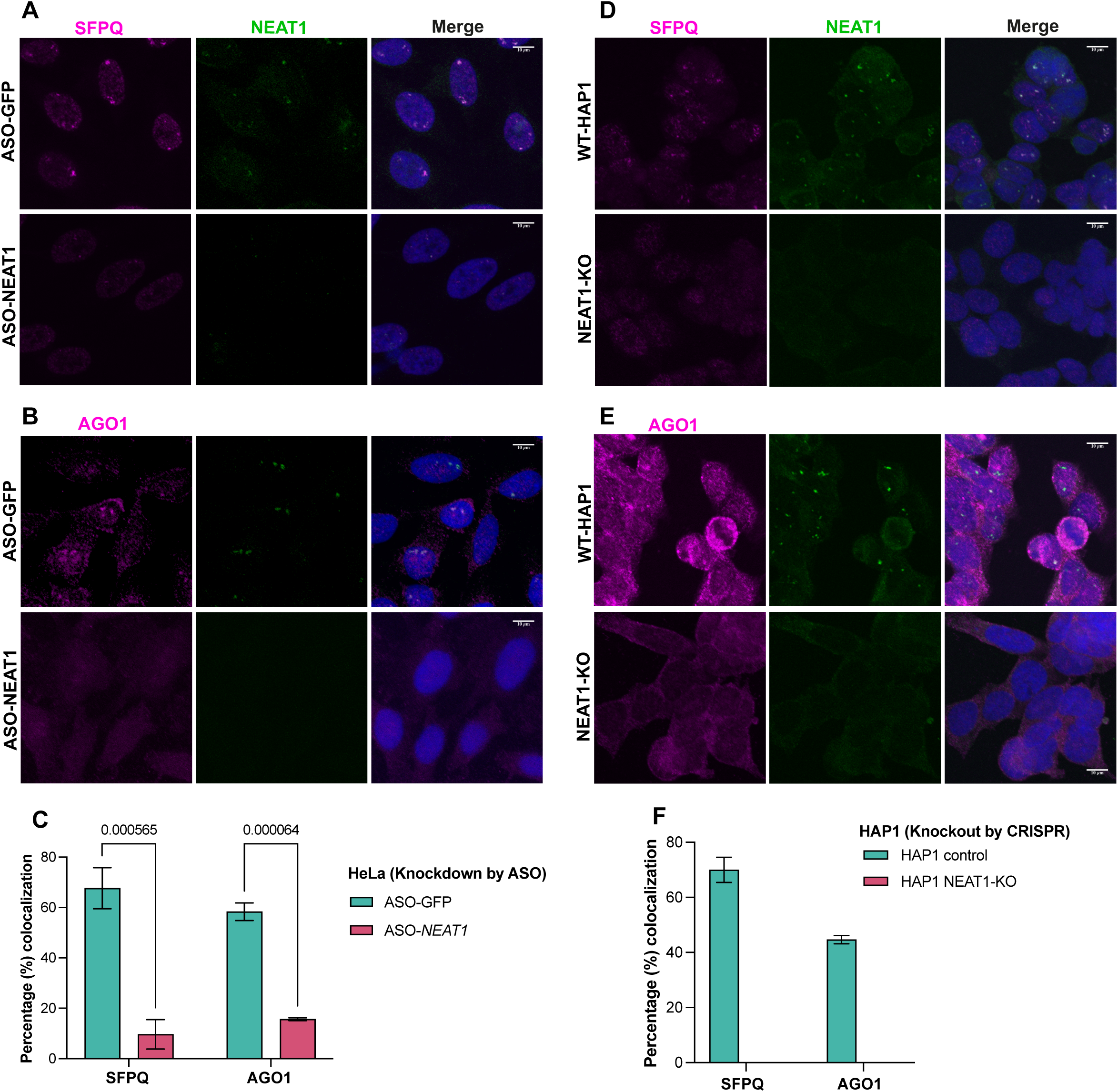
Depletion of *NEAT1* effect AGO1 localization. Transient depletion of *NEAT1* by antisense oligo (ASO-*NEAT1*) affects AGO1 localization in the nucleus. GFP antisense oligo (ASO-GFP) was used as negative control. (**A**) SFPQ and *NEAT1* RNA-FISH colocalization in the control (ASO-GFP) and ASO-*NEAT1* condition. (**B**) AGO1 and *NEAT1* RNA-FISH colocalization in the control (ASO-GFP) and ASO-*NEAT1* condition. (**A-B**) In *NEAT1* depleted cells both AGO1 and SFPQ signals disappeared from paraspeckles. (**C**) Quantification of SFPQ and AGO1 signals within *NEAT1* paraspeckles in the control (ASO-GFP) and ASO-*NEAT1* condition. (**D-F**) Similar to (**A-C**) *NEAT1* knockout (KO) HAP1 cell lines displayed a diffuse (**D**) SFPQ and (**E**) AGO1 signals compared to WT-HAP1. (**F**) Quantification of SFPQ and AGO1 signals within paraspeckles in *NEAT1-*KO cells.

### AGO1 is required for paraspeckle integrity

Our previously reported AGO1 ChIP-seq data revealed significant enrichment at the promoter of *NEAT1* gene (Figure 5A), indicating that *NEAT1* locus may be a direct target of AGO1. Notably, AGO1 ChIP-peaks at *NEAT1* promoter also overlapped with binding sites of two transcription factors (HSF1 and ATF2) that are known to be involved in *NEAT1* induction(Lellahi et al., 2018; Wang et al., 2018) (Figure S1). Therefore, we measured the expression of both *NEAT1_1* and *NEAT1_2* in AGO1 knockdown cells. Interestingly, we observed up-regulation of both *NEAT1* transcripts (*NEAT1_1* and *NEAT1_2*) upon AGO1 depletion (Figure 5B). The expression of *NEAT1* is an essential early step in paraspeckle formation, thus we asked whether AGO1 is involved in paraspeckle assembly. We examined the paraspeckle integrity after AGO1 knockdown by analyzing co-localization of *NEAT1* with SFPQ an essential paraspeckle protein component. Despite increased *NEAT1* RNA-FISH signal, we observed a diffuse SFPQ signal in >60% of siAGO1 cells, affecting paraspeckle integrity (Figure 5C-D). Furthermore, by ChIRP-western blot we checked the effect of AGO1 depletion on the interaction of other key paraspeckle proteins with *NEAT1.* In AGO1 depleted cells, immunoblot analyses of *NEAT1* associated proteins showed a significant reduction in the level of crucial paraspeckle components (NONO, HNRNPK, and SFPQ) (Figure 5E-F). However, the total level of these paraspeckle proteins (NONO, SFPQ, HNRNPK, and PSPC1) remained unchanged in AGO1 knockdown cells (Figure 5G). Similarly, the ChIRP experiment demonstrated comparable levels of *NEAT1* enrichment under both control and AGO1 knockdown conditions (Figure S2).

**Figure 5.**
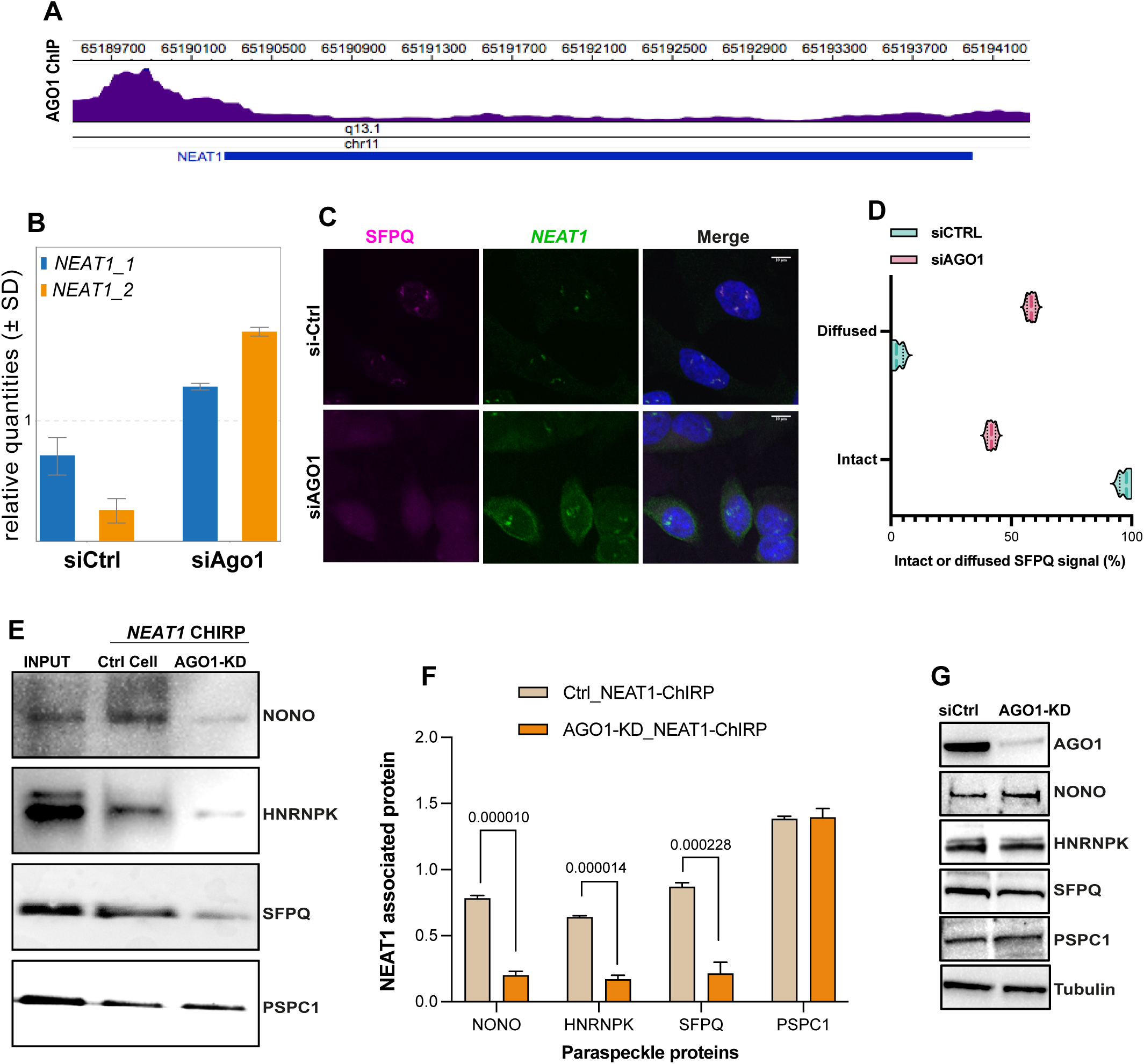
AGO1 knockdown impacts paraspeckles formation, *NEAT1* expression, and its interaction with other PSPs. (**A**) Enrichment of AGO1 ChIP-seq signal (purple) on the promoter locus (chr11) of *NEAT1* lncRNA. (**B**) RT-qPCR analysis of *NEAT1_1* and *NEAT1_2* expression in siCtrl and siAGO1 cells. Values were normalized relative to the geometric mean of 18S rRNA, GAPDH, and Actin-B mRNA expression levels. Error bars indicate mean ± sd. (**C**) Examining the effects of AGO1 depletion on paraspeckle formation. Paraspeckle integrity was examined by SFPQ localization to *NEAT1* paraspeckles using a combination of RNA-FISH and immunofluorescence. (**D**) Quantification of the number of cells having paraspeckles with intact or diffuse SFPQ signal as shown in section **C**. The graph represents mean (± s.e.m) of three technical replicates. We scored paraspeckle morphology in 670 cells (for the combined replicates) of siCtrl and siAGO1 cells. (**E**) Western blot for some essential paraspeckle proteins (NONO, SFPQ, HNRNPK and PSPC1) retrieved with *NEAT1* ChIRP in control and AGO1 knockdown cells. AGO1 knockdown decreases the interaction of NONO, SFPQ, and HNRNPK with *NEAT1*. (**F**) Quantification *NEAT1*-ChIRP western-blot data (n=3) as shown in **E**. (**G**) The total level of paraspeckle proteins (NONO, SFPQ, HNRNPK and PSPC1) was not changed upon AGO1 depletion. Tubulin was used as a loading control.

### Loss of *NEAT1* impacts active chromatin architecture

*NEAT1* was reported to be enriched on active chromatin by CHART-seq in human cells (West et al., 2014). We observed that around 94% of *NEAT1* CHART-seq peaks overlapped with AGO1 binding (ChIP-seq) at active sites (Figure 6A and Figure S3). This finding suggests that AGO1 may work in concert with *NEAT1* paraspeckles to regulate genome architecture. To test this hypothesis, we examined how the loss of *NEAT1*-paraspeckles affects chromatin organization using Hi-C in wild-type (WT) and *NEAT1*-KO HAP1 cell lines. *NEAT1*-KO cell lines from two clones showing similar cell-cycle profiles compared to WT HAP1 cell line replicates (Figure S4) were processed for further analysis. We determined significant interaction matrices with the HiC-Pro pipeline(Servant et al., 2015) (Figure S5), differential interactions by diffHiC (Lun and Smyth, 2015), and (A/B) compartment analysis by principal components. The HiC from two independent clones of *NEAT1*-KO cell lines display significant correlation (Figure S6), indicating that there is no clonal effect. In *NEAT1*-KO cells, differential Hi-C analysis revealed 5,674 bins with significantly decreased interactions (FDR < 0.05, logFC > 1.2) and 390 bins with increased interactions. Furthermore, comparing genome-wide compartments between *NEAT1*-KO and WT-HAP1 cells identified significant changes in A/B chromatin compartments (Figure 6B-C). Notably, we observed switching of A to B compartments (Fisher’s test, FDR < 0.05) across multiple regions in most chromosomes (Figure 6B-C). As an example, the PC1 values illustrate the B to A compartment switching on chromosome 16 between WT and *NEAT1*-KO HAP1 cells (Figure 6C). The changes in chromatin compartmentalization were further highlighted by the weak correlation of eigenvectors between WT and *NEAT1*-KO cells, particularly on chromosomes 4, 9, 21, and 22 (Figure 6D). The observed compartment switching upon *NEAT1* knockout, mirroring the effects of AGO1 depletion in our previous work(Shuaib et al., 2019), underscoring their shared role in shaping the chromatin landscape.

**Figure 6.**
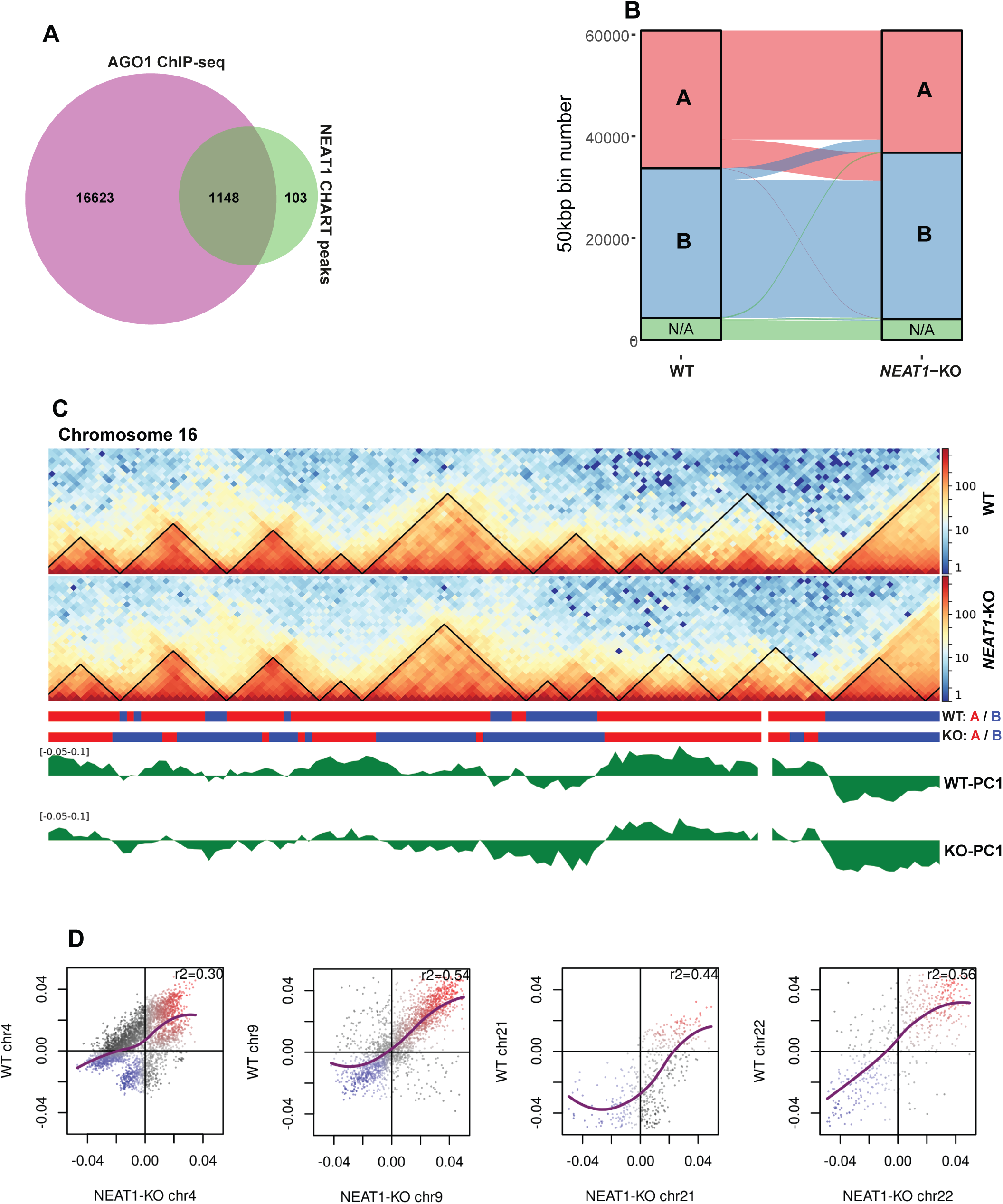
Loss of *NEAT1* affects chromatin compartmentalization. (**A**) Venn diagram shows overlapping of AGO1 ChIP-seq peaks with *NEAT1* CHART-seq data (Data accession# GSM1411207 and GSM1411208). (**B**) Alluvial plot shows A and B compartment profile between WT HAP1 and *NEAT1*-KO cells. Compartments switching from A-type to B-type in *NEAT1*-KO cells are shown. (**C**) Representation of TADs and compartments in Chromosome-16 (locus 50,000,000-60,000,000) identified in WT (top) and *NEAT1* knockout (bottom) HAP1 cells. The A-type compartments are colored red and B-type compartments are shown in blue. PC1 values of chromosome 16 used to define A/B chromatin compartments, positive values represent A-type and negative values represent B-type compartments in WT-HAP1 (top) and *NEAT1* knockout (bottom) HAP1 cells. (**D**) First eigenvector correlation for chromosome 4, 9, 21 and 22 between WT-HAP1 and *NEAT1* knockout conditions are shown.

## Discussion

Early studies in fission yeast and plants established a role for nuclear RNAi components, particularly Argonaute (AGO) proteins, in the epigenetic regulation of heterochromatin formation (Volpe et al., 2002; Zilberman et al., 2003). Surprisingly, subsequent research in animal cells revealed that AGO proteins are enriched at transcriptionally active chromatin sites (Allo et al., 2014; Cernilogar et al., 2011; Huang et al., 2013; Shuaib et al., 2019), raising new questions about their nuclear function. In this work, we demonstrate that human nuclear AGO1 is localized to paraspeckles and functions in the assembly of these sub-nuclear structures. Paraspeckles are nuclear subdomains that contain various RNAs and RNA-binding proteins (RBPs). *NEAT1* lncRNA is the key organizer of paraspeckles and is required for maintaining its structural integrity (Clemson et al., 2009; Fox et al., 2002; Sasaki et al., 2009). Through RIP-seq analysis, we identified *NEAT1* as a major nuclear partner of AGO1, along with other RNA species, including lncRNAs, mRNAs, and short RNAs. While we do not exclude binding to short RNAs, because of the preferential association of AGO1 with *NEAT1* lncRNA in the nucleus, we focused on exploring the functional link of nuclear AGO1 with paraspeckles. Our findings establish AGO1 as a novel component of paraspeckles, supported by several key observations: AGO1 paraspeckle localization, its direct interaction with *NEAT1*, the loss of AGO1 localization upon *NEAT1* depletion, its association with essential paraspeckle proteins, and the effects of AGO1 knockdown on *NEAT1* expression and paraspeckle integrity.

The evidence that AGO1 localizes to paraspeckles and interacts with key components, including *NEAT1* lncRNA and paraspeckle proteins (PSPs), strongly suggests a role for AGO1 in paraspeckle formation. Paraspeckles formation involves two main steps: (1) the transcription of both *NEAT1* isoforms (*NEAT1_1* and *NEAT1_2*) and (2) the assembly of essential paraspeckle proteins around *NEAT1_2* (Chujo et al., 2017; Clemson et al., 2009; Mao et al., 2011; Naganuma et al., 2012; Sasaki et al., 2009). Our data show that AGO1 depletion leads to hyper-expression of both *NEAT1* isoforms and reduces the interaction of key paraspeckle proteins, such as SFPQ and NONO, with *NEAT1* lncRNA. This supports a role for AGO1 in regulating both *NEAT1* expression and paraspeckle assembly. The *de novo* assembly of paraspeckles is initiated in the vicinity of *NEAT1* transcription site (Mao et al., 2011). *NEAT1_2* is the essential structural unit of paraspeckles(Sasaki et al., 2009), which is produced from alternative 3′-end processing of *NEAT1_1 (Naganuma et al., 2012)*. Interestingly, HNRNPK, a protein that interacts with AGO1, has been shown to regulate *NEAT1_2* synthesis by modulating *NEAT1_1*’s 3′-end processing (Naganuma et al., 2012). The primary transcription factors involved in stress-mediated *NEAT1* induction include heat shock transcription factor 1 (HSF1) (Lellahi et al., 2018) and activating transcription factor 2 (ATF2) (Wang et al., 2018). Interestingly, AGO1 peaks at *NEAT1* promoter overlapped with binding sites of these transcription factors. These findings collectively suggest that nuclear AGO1 plays an essential role in the transcriptional and post-transcriptional processes that are critical for paraspeckle formation.

Though the exact function of paraspeckles is not fully explored, their dynamic assembly in stress-response is believed to regulate gene expression by distinct mechanisms (Hirose et al., 2014; Imamura et al., 2014; Prasanth et al., 2005). Recent work, using a series of *NEAT1* deletion mutants, discovered a functional domain in the middle region of *NEAT1_2*, which is essential for the final assembly of paraspeckles(Yamazaki et al., 2018). The binding of SFPQ and NONO proteins to the functional middle domain of *NEAT1_2* followed by the recruitment of additional PSPs induces condensation mediated paraspeckle assembly (Yamazaki et al., 2018). Many key paraspeckle proteins (like NONO, FUS, and RBM14) contain intrinsically disordered domains that can facilitate liquid–liquid phase separation (Fox et al., 2018; Yamazaki et al., 2018). Intriguingly, we also found some predicted intrinsically disorder regions in the AGO1 protein (Figure S7) by using the online server MetaDisorder (Kozlowski and Bujnicki, 2012). Also, our data display strong enrichment of AGO1 peaks on the middle domain of *NEAT1_2*. Moreover, both AGO1 and *NEAT1* lncRNA exhibited overlapping binding at active chromatin sites. This overlap suggests a potential role for the AGO1-*NEAT1* lncRNA interaction in modulating the aggregation states of genomic elements, potentially influencing nuclear architecture.

Interestingly, depletion of *NEAT1* induces changes in chromatin active compartments, an effect like what we observed upon AGO1 depletion. This parallel suggests a coordinated role for AGO1 and *NEAT1* lncRNA in shaping the chromatin landscape. Given that AGO1 interacts with chromatin regulators (Fallatah et al., 2021; Shuaib et al., 2019) and transcriptional machinery (Huang et al., 2013), its association with *NEAT1* likely serves dual functions: facilitating paraspeckle formation and modulating chromatin architecture. The fact that *NEAT1* depletion mirrors AGO1 depletion in terms of chromatin compartment switching reinforces the idea that paraspeckles, guided by *NEAT1*, might be involved in regulating chromatin states. This link between paraspeckles and chromatin structure offers intriguing directions for future research, particularly in exploring how nuclear bodies contribute to the regulation of genome architecture.

## Methods

### Cell Lines

HepG2 (ATCC, HB-8065, male), HeLa (ATCC, CCL-2, female), HEK-293 (ATCC, CRL-3216, female), HAP1 (Horizon) cells were purchased before experiments and thus considered authenticated.

### Cell culture and transfection

HepG2, HeLa and HEK-293 cells were cultured in EMEM medium (Sigma, Cat # M0643) supplemented with 10% FBS (Invitrogen, Cat # 26140-079), and HAP1 cells were grown in Iscove’s Modified Dulbecco’s Medium (IMDM, GIBCO, Cat.No. 12440-053) with 10% FBS, 1% penicillin and streptomycin (Euroclone, Cat # ECB3001D) was added to culture media. Cell cultures were maintained at 37°C and 5% CO2. AGO1 knockdown experiments were performed using a pool of four siRNAs, as described previously (Shuaib et al., 2019). The phosphothioate-converted antisense oligonucleotides (ASO) against *NEAT1* or negative control GFP (See below the sequence information) were used as describe previously (Sasaki et al., 2009).

| Anti-sense Oligo | Sequence (5' - 3') |
| --- | --- |
| **GFP-ASO** | mU*mC*mA*mC*mC*T*T*C*A*C*C*C*T*C*T*mC*mC*mA*mC*mU |
| **NEAT1-ASO-1** | mC*mC*mC*mU*mC*T*A*G*T*C*T*T* G*G*C*mU*mC*mA*mU*mU |
| **NEAT1-ASO-2** | mC*mC*mC*mU*mU*C*T*C*C*T*A*G* T*A*A*mU*mC*mU*mG*mC |
| **Description**<br>* for Phosphorothioate backbone<br>N represent deoxynucleotide<br>mN for represent 2'-O-methylribonucleotide |  |

The *NEAT1* knockout HAP1 cell lines were previously generated (Yamazaki and Hirose, 2021) by deleting the whole *NEAT1* gene locus using two sgRNAs targeting the 5’- and 3’-ends of *NEAT1* (sgRNA#1: TCCCTCCCTGTCGCTAACTC, sgRNA#2: GCAAAACCTGAGTGCGGCCA). Two independent clones were selected for downstream analysis.

### Antibodies

Mouse monoclonal anti-AGO1 (a kind gift from Mikiko Siomi), anti-AGO1 (clone 2A7, Wako), anti-SFPQ (RN014MW, Ribonomics), anti-HNRNPK (SAB4501424, Sigma), and anti-TLS/FUS (ab23439, Abcam), and anti-PSPC1 (ab104238, Abcam), β-Tubulin antibody (#2146, Cell Signaling), and Histone H3 antibody (#9715, Cell Signaling), goat anti-NONO /p54NRB (EB07246) were used as primary antibody for western blotting and or immunofluorescence. Alexa flour 647 Goat anti-mouse IgG (H+L) (ab150115, Abcam) and Alexa flour 488 Donkey anti-mouse IgG (H+L) (Molecular probe, A21202) were used as secondary antibody for immunofluorescence.

### Chromatin bound AGO1 RIP-seq analysis

The AGO1 RIP-seq analysis for both 100 and 400 fragment sizes was carried out as described previously (Shuaib et al., 2019). For UV cross-linking IP combined with qRT-PCR, cells were cross-linked on ice with 254 nM UV-C at 0.3 J/cm2, and the rest of the experiment was performed as described previously (Sasaki et al., 2009).

### RT-qPCR

Total RNA was extracted with TRI reagent (Sigma, Cat # T9424) and reverse transcribed with the QuantiTect reverse transcription kit (Qiagen, Cat # 205311). The resulting cDNA was then amplified via quantitative PCR on CFX96 Real-Time PCR Machine (Bio Rad) using the SYBR Select Mastermix (Applied Biosystems). For relative quantification of RNA levels, the geometric mean of 18S rRNA, GAPDH, and Actin-B mRNA was used as normalization reference. Human AGO1-specific primer was purchased from Qiagen (Cat# QT00006370) and reference genes (18S, GAPDH, and ActB) primers were sourced from geNorm Syber green kit (Primerdesign Cat# ge-SY-12).

For RT-qPCR analysis of human *NEAT1_1* and *NEAT1_2* expression two set of primers were used. The set-1 recognizes only *NEAT1_1* while set-2 can amplify both *NEAT1_1* and *NEAT1_2*.

Set-1:

Fw: TTGTTCCAGAGCCCATGAT

Rv: TGAAAACCTTTACCCCAGGA

Set-2:

Fw: GATCTTTTCCACCCCAAGAGTACATAA

Rv: CTCACACAAACACAGATTCCACAAC

### Fluorescent *in situ* hybridization (RNA-FISH) and Immunocytochemistry

The probes for *NEAT1* lncRNA were synthesized by *in-vitro* transcription of antisense RNA using linearized plasmids containing a *NEAT1* fragment (+1 to +1,000). The experiments were performed as described previously (Naganuma et al., 2012).

### Tandem affinity purification and mass spectrometry

The soluble nuclear extracts and chromatin bound extracts were prepared from stable HEK-293 cell lines expressing either AGO1 or AGO2 proteins fused to C-terminal FLAG and HA epitope tags (tagged protein are denoted as e-AGO1 and e-AGO2). The stable cells lines were produced (Shuaib et al., 2019) using retroviral vector (pOZ-FH-C-puro, Addgene plasmid # 32516) (Kumar et al., 2009). The e-AGO1 and e-AGO2 protein complexes were purified by double-immunoaffinity purification procedure from nuclear soluble and chromatin fraction with anti-Flag M2 antibody-conjugated agarose (Sigma), followed by anti-HA purification. Interacting protein partners of purified complexes (AGO1-com and AGO2-com) were identified by mass spectrometer (Bioscience Core laboratory, KAUST). The interactions with specific partners were further confirmed by western blotting.

### *NEAT1* ChIRP and western blotting

*NEAT1* ChIRP followed by protein isolation was performed as described previously (Chu et al., 2015). Briefly, 500 million cells (per ChIRP reaction) were cross-linked with formaldehyde, target RNA was retrieved with Magna ChIRP *NEAT1* probes set (even, odd) (Cat. # 03-308) or Magna ChIRP negative control probe (LacZ, part # CS216572), and proteins were eluted with biotin. The enriched proteins were used for silver staining and western blotting.

### Hi-C libraries preparation and data analysis

Hi-C libraries were prepared from 25 million HAP1 cells (WT and *NEAT1*-KO) as described previously described (Shuaib et al., 2019). Hi-C data analysis including differential interactions, TAD analysis, and compartment identification were performed(Shuaib et al., 2019). We obtained around 690 million valid pairs, ∼358 million for WT-HAP1 and ∼331 million for *NEAT1*-KO HAP1 cell lines.

## Data availability

The sequencing data such as AGO1 (chromatin bound) RIP-seq and AGO1 ChIP-seq have been previously (Shuaib et al., 2019) deposited at the Sequence Read Archive (SRA, http://www.ncbi.nlm.nih.gov/sra/), with accession number SRA: PRJNA398595. HAP1 Hi-C data for WT and *NEAT1*-KO conditions is available under accession number: PRJNA1168304.

## Acknowledgments

We are grateful to Heno Hwang (scientific illustrator) at KAUST for his help in drawing the model; KAUST Bioscience Core Lab for providing sequencing facility. This work was supported by baseline fund from the King Abdullah University of Science and Technology (KAUST) to V.O. The grants from the MEXT of Japan to TH (21H05276 and 24K21933) and JST CREST to TH (JPMJCR20E6). The grant for Joint Research Program of IGM, Hokkaido University.

