## Supplemental Figures for "AGO1 interacts with *NEAT1* lncRNA and impacts nuclear compartments"

**Supplementary Figure Legends:**

**Figure S1.** Enrichment of AGO1, HSF1, and ATF2 ChIP-seq signal on the promoter locus (chr11) of *NEAT1* lncRNA.

**Figure S2. *NEAT1* ChIRP experiment. (A)** Sketch showing *NEAT1* ChIRP procedure. **(B)** *NEAT1* ChIRP was performed in siCtrl, siAGO1, and LacZ probes were used as negative control. Isolated RNAs were analyzed by qRT-PCR using specific primers for *NEAT1* and GAPDH (negative target). Bar chart shows similar level of *NEAT1* lncRNA retrieval (%) in both conditions (n=3, paired t-test p-value =ns).

**Figure S3. AGO1 ChIP-seq and *NEAT1* CHART-seq signals.** Some representative regions showing overlap of AGO1 ChIP-peaks and *NEAT1* CHART-peaks.

**Figure S4. Cell cycle analysis**. Cells were analyzed by flow cytometry after PI staining to measure DNA content. Flow cytometry analysis display similar cell cycle profiles for **(A)** wildtype HAP1 cell lines and **(B)** *NEAT1*-KO HAP1 cell lines from two independent clones.

**Figure S5. Statistics and quality control of HiC libraries. (A)** The mapping and HiC library statistics for both replicates of wildtype HAP1 cell lines (WT1 and WT2) and *NEAT1*-KO HAP1 cell lines (KO_1 and KO_2). Genome-wide normalized HiC interaction heatmaps at 1-Mb resolution are shown in **(B)** wildtype HAP1 **(C)** *NEAT1*-KO HAP1.

**Figure S6.** Spearman correlation of HiC libraries in wildtype HAP1 cell lines (WT1 and WT2) and *NEAT1*-KO HAP1 cell lines (KO_1 and KO_2).

**Figure S7. AGO1 protein contains intrinsically disordered residues**. Prediction of disordered amino acid residues in the AGO1 protein by using MetaDisorder tool. The threshold is shown by red line at 0.5 on the y-axis for disordered/ordered residues. The predicted disordered residues are having score above this red line.

### Figure S1

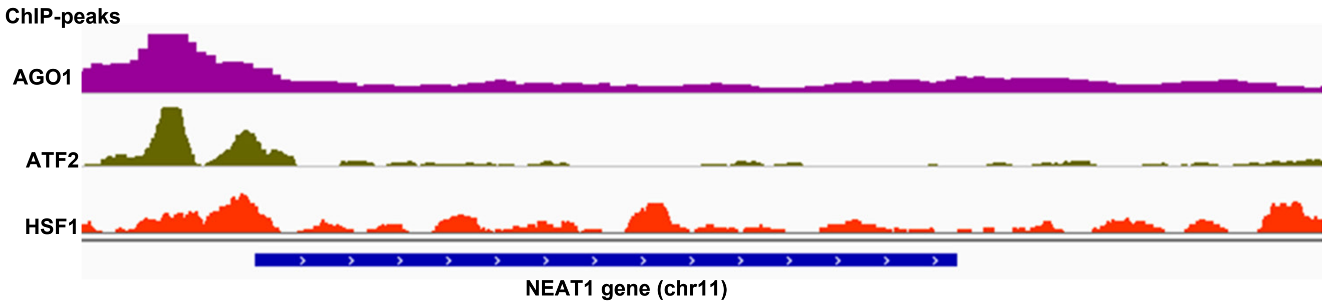

#### Figure S2

A

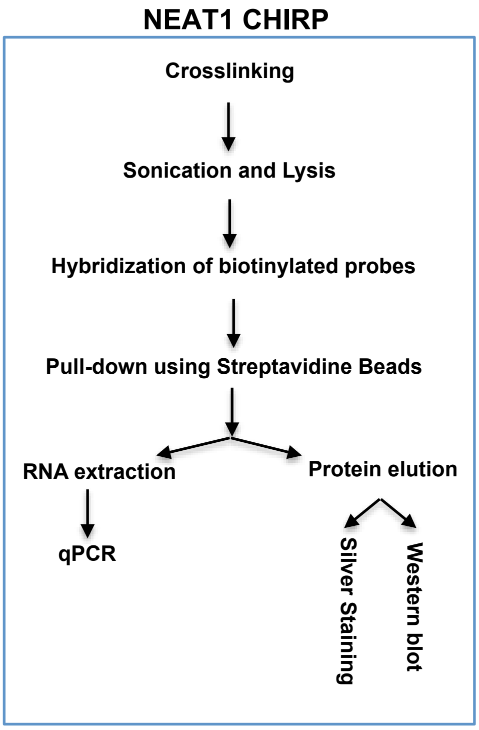
NEAT1 CHIRP
Crosslinking
Sonication and Lysis
Hybridization of biotinylated probes
Pull-down using Streptavidine Beads
RNA extraction
Protein elution
qPCR
Silver Staining
Western blot

B

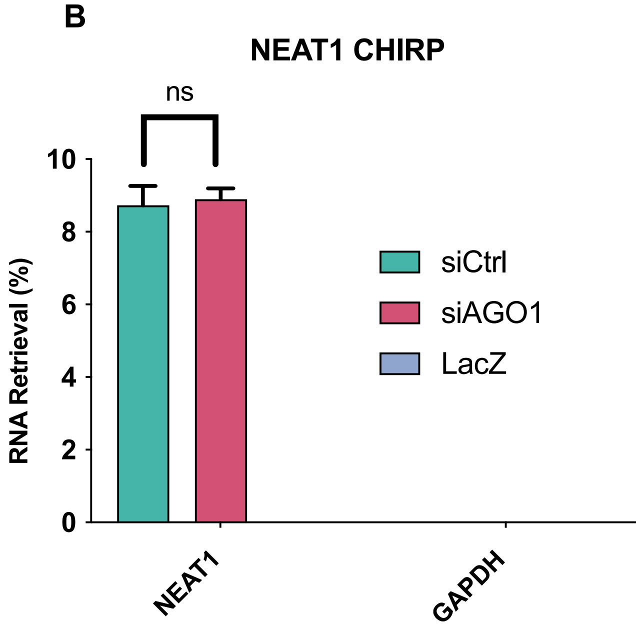
NEAT1 CHIRP
ns
RNA Retrieval (%)
0
2
4
6
8
10
siCtrl
siAGO1
LacZ
NEAT1
GAPDH

### Figure S3

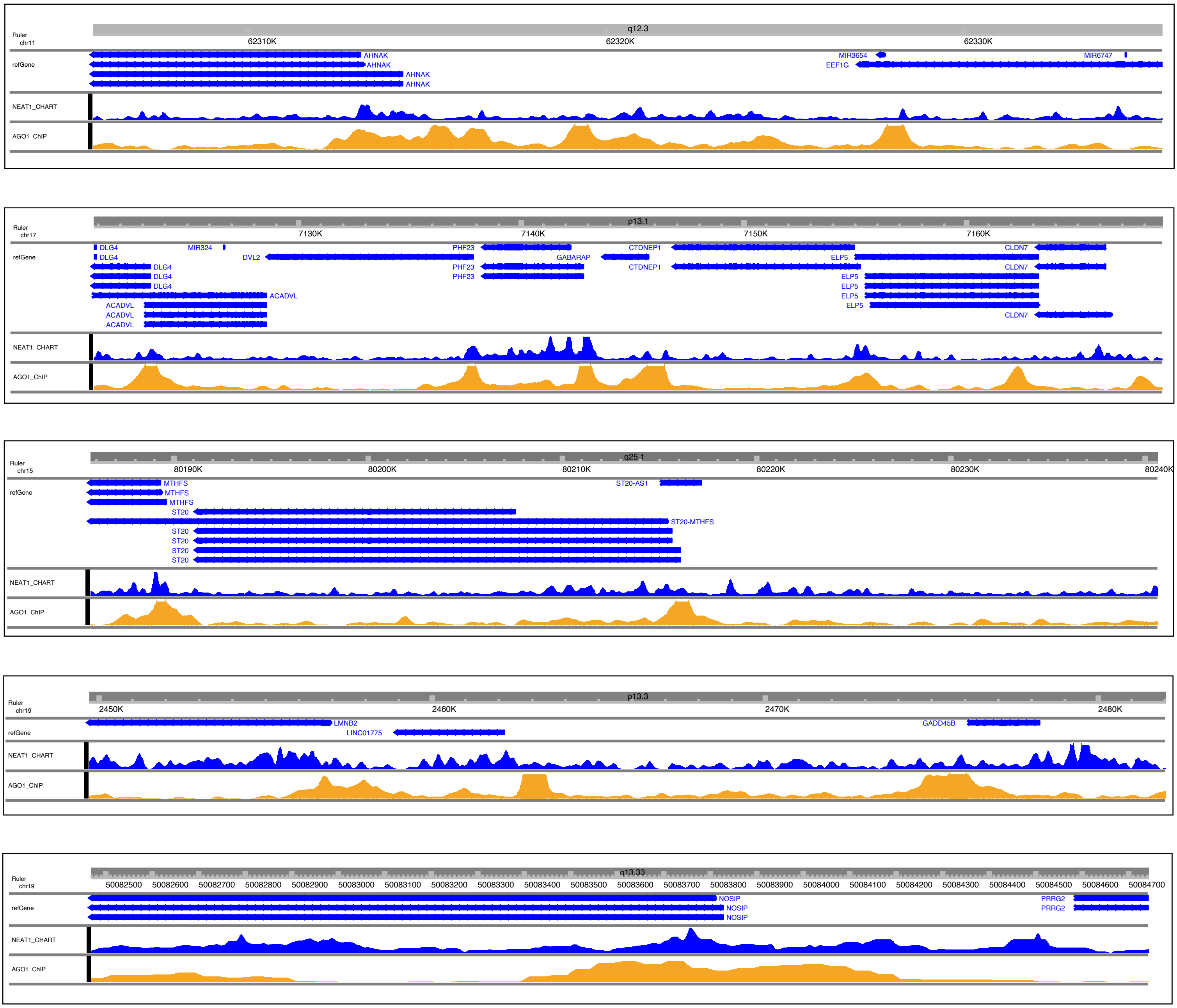

Ruler
chr11
q12.3
62310K
62320K
62330K
refGene
AHNAK
AHNAK
AHNAK
AHNAK
MIR3654
EEF1G
MIR6747
NEAT1_CHART
AGO1_ChIP
Ruler
chr17
p13.1
7130K
7140K
7150K
7160K
refGene
DLG4
DLG4
DLG4
DLG4
DLG4
MIR324
DVL2
ACADVL
ACADVL
ACADVL
ACADVL
PHF23
PHF23
PHF23
GABARAP
CTDNEP1
CTDNEP1
ELP5
ELP5
ELP5
ELP5
ELP5
CLDN7
CLDN7
CLDN7
NEAT1_CHART
AGO1_ChIP
Ruler
chr15
q25.1
80190K
80200K
80210K
80220K
80230K
80240K
refGene
MTHFS
MTHFS
MTHFS
ST20
ST20-MTHFS
ST20
ST20
ST20
ST20
ST20-AS1
NEAT1_CHART
AGO1_ChIP
Ruler
chr19
p13.3
2450K
2460K
2470K
2480K
refGene
LMNB2
LINC01775
GADD45B
NEAT1_CHART
AGO1_ChIP
Ruler
chr19
q13.33
50082500
50082600
50082700
50082800
50082900
50083000
50083100
50083200
50083300
50083400
50083500
50083600
50083700
50083800
50083900
50084000
50084100
50084200
50084300
50084400
50084500
50084600
50084700
refGene
NOSIP
NOSIP
NOSIP
PRRG2
PRRG2
NEAT1_CHART
AGO1_ChIP

### Figure S4

## A

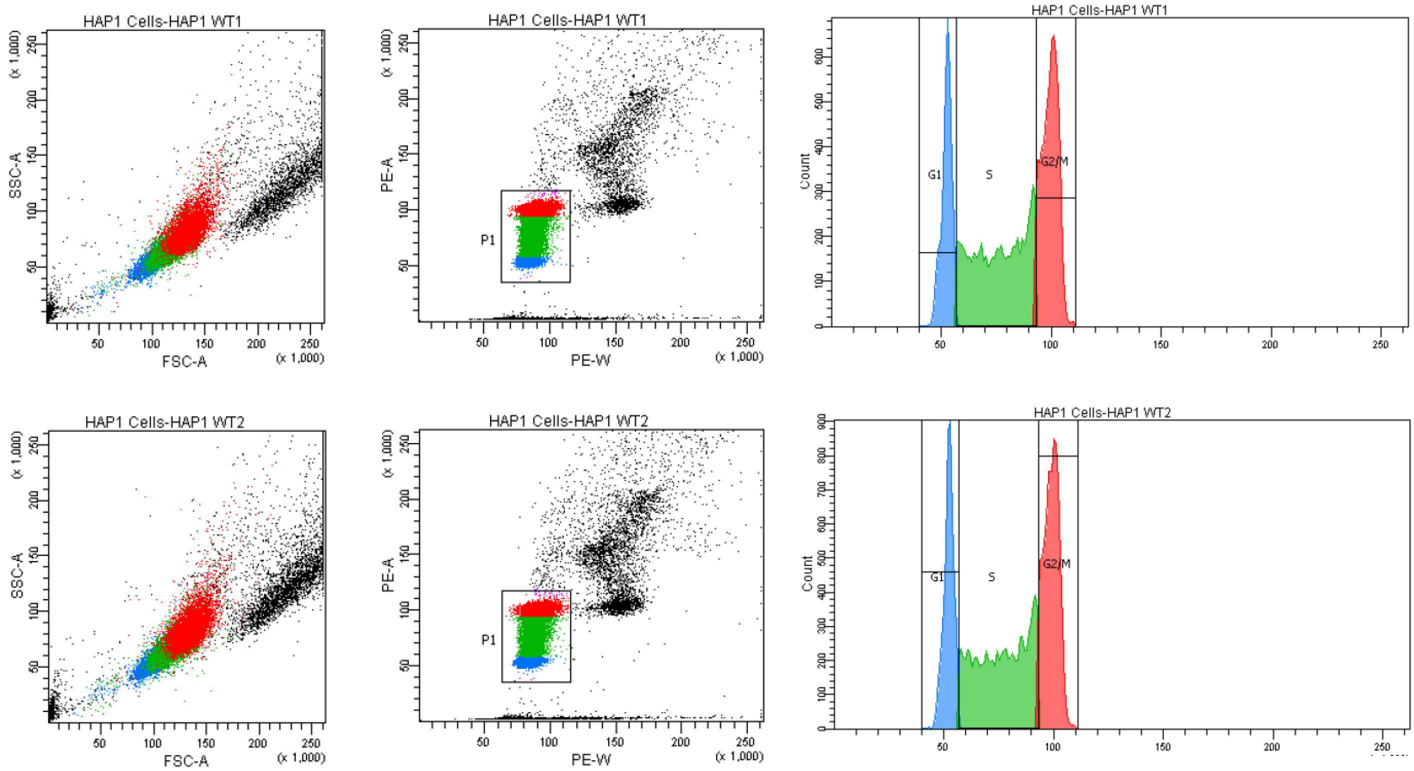
HAP1 Cells-HAP1 WT1
SSC-A
FSC-A
(x 1,000)
PE-A
PE-W
P1
Count
G1
S
G2/M
HAP1 Cells-HAP1 WT2

## B

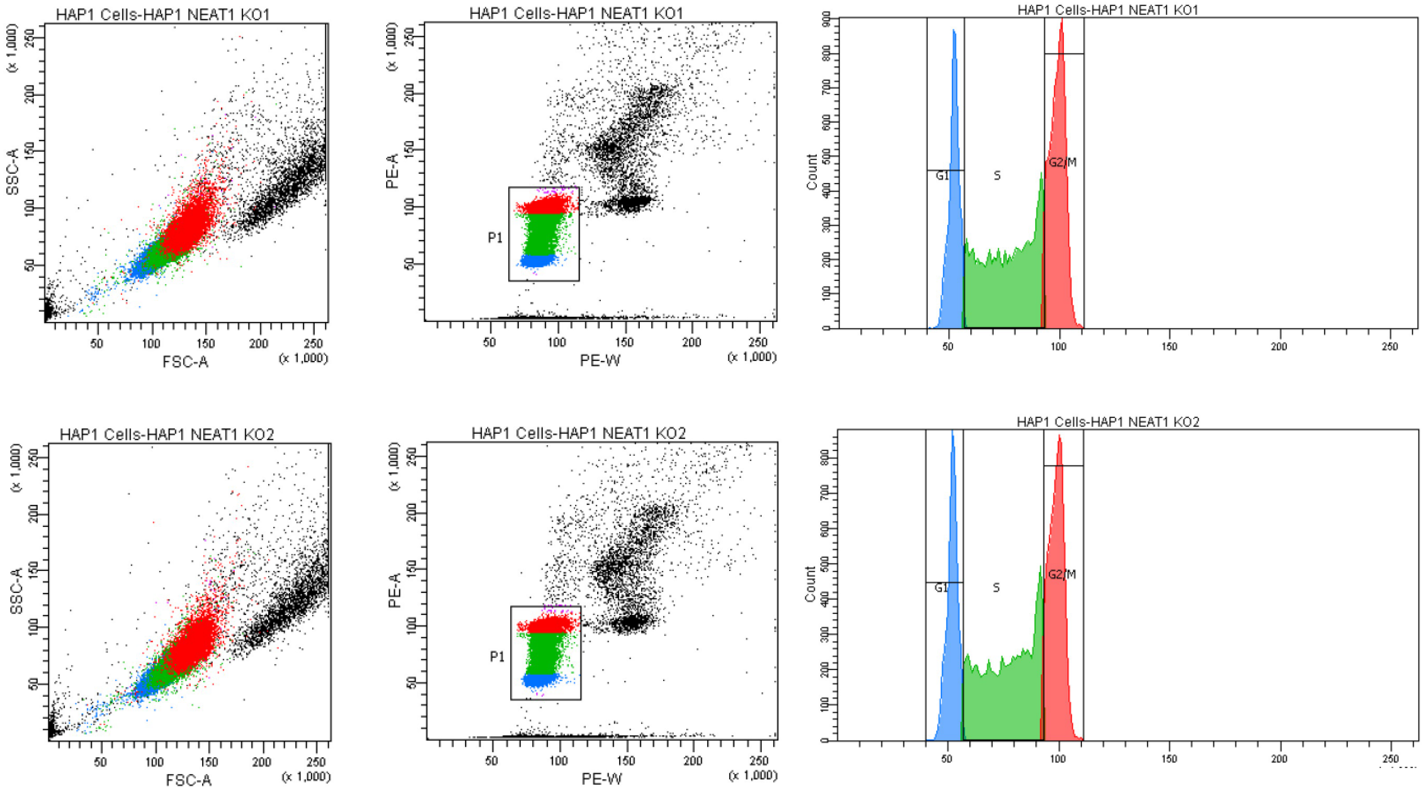
HAP1 Cells-HAP1 NEAT1 KO1
SSC-A
FSC-A
(x 1,000)
PE-A
PE-W
P1
Count
G1
S
G2/M
HAP1 Cells-HAP1 NEAT1 KO2

### Figure S5

## A

| HiC Statistics ( Replicates) | | | | |
|---|---|---|---|---|
| | HAP1-WT1 | HAP1-WT2 | NEAT1_KO1 | NEAT1_KO2 |
| Raw (pairs) | 346271757 | 448756409 | 351303056 | 498089738 |
| Trimmed q20 (pairs) | 297840334 | 378102768 | 315010026 | 427159192 |
| Ratio Trimmed/Raw | 0.860134643 | 0.84255681 | 0.89669025 | 0.85759485 |
| Mapped uniquely (pairs) | 249699653 | 318756563 | 268521844 | 360852317 |
| Mapping efficiency | 83.83674892 | 84.304213 | 85.2423167 | 84.477245 |
| Self-circle | 3199049 | 4725117 | 3626846 | 4859366 |
| Dangling-end | 5493400 | 6163629 | 8496563 | 11655870 |
| Error | 674341 | 1680870 | 1695901 | 2237618 |
| Extra dangling-end | 50323541 | 66122073 | 63064421 | 83778610 |
| Too short | 2465816 | 4219702 | 3926096 | 5324403 |
| Too large | 238734 | 277367 | 276880 | 361017 |
| Duplicated | 55531156 | 62518716 | 28324170 | 81496222 |
| Random breaks | 4917687 | 6402748 | 8143701 | 11782178 |
| Total valid | 144193401 | 187406098 | 165688097 | 192818732 |
| Total valid (% relative to Raw) | 41.64 | 41.76 | 47.16 | 38.71 |
| Total valid (% relative to Trimmed) | 48.41 | 49.56 | 52.60 | 45.14 |
| Total valid (% relative to Mapped uniquely) | 57.75 | 58.79 | 61.70 | 53.43 |

| HiC Statistics (Combined Samples) | | |
|---|---|---|
| | HAP1-WT | NEAT1_KO |
| Raw (pairs) | 795028166 | 849392794 |
| Trimmed q20 (pairs) | 675943102 | 742169218 |
| Ratio Trimmed/Raw | 0.85021277 | 0.87376444 |
| Mapped uniquely (pairs) | 568456216 | 629374161 |
| Mapping efficiency | 84.0982346 | 84.8019758 |
| Self-circle | 7924166 | 8486212 |
| Dangling-end | 11657029 | 20152433 |
| Error | 2355211 | 3933519 |
| Extra dangling-end | 116445614 | 146843031 |
| Too short | 6685518 | 9250499 |
| Too large | 516101 | 637897 |
| Duplicated | 118049872 | 109820392 |
| Random breaks | 11320435 | 19925879 |
| Total valid | 331599499 | 358506829 |
| Total valid (% relative to Raw) | 41.71 | 42.21 |
| Total valid (% relative to Trimmed) | 49.06 | 48.31 |
| Total valid (% relative to Mapped uniquely) | 58.33 | 56.96 |

## B

WT-HAP1

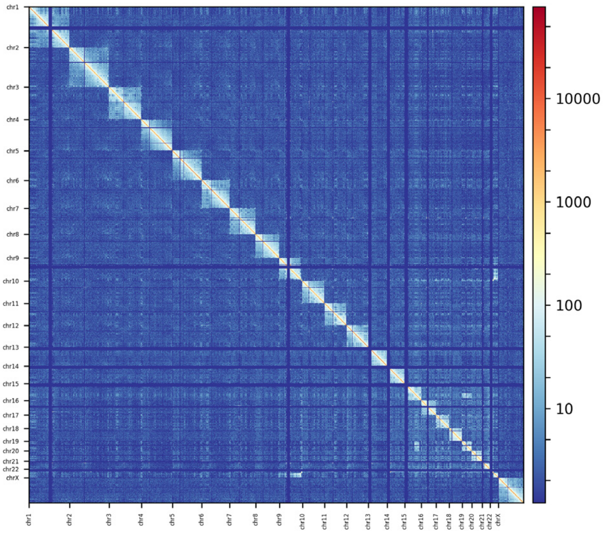

## C

NEAT1-KO-HAP1

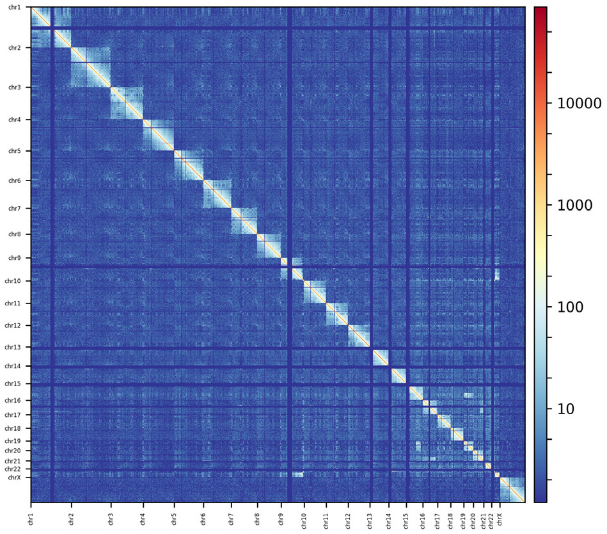

**Figure S6**

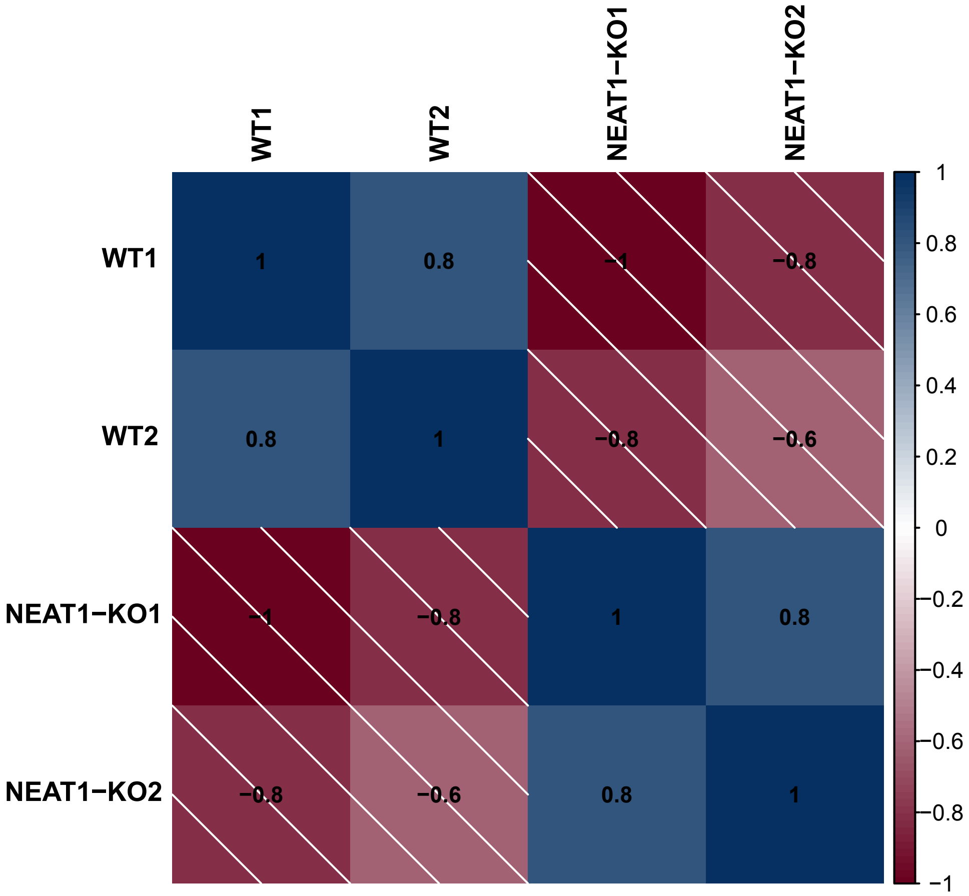

WT1
WT2
NEAT1−KO1
NEAT1−KO2
WT1
WT2
NEAT1−KO1
NEAT1−KO2
1
0.8
−1
−0.8
0.8
1
−0.8
−0.6
−1
−0.8
1
0.8
−0.8
−0.6
0.8
1
1
0.8
0.6
0.4
0.2
0
−0.2
−0.4
−0.6
−0.8
−1

#### Figure S7

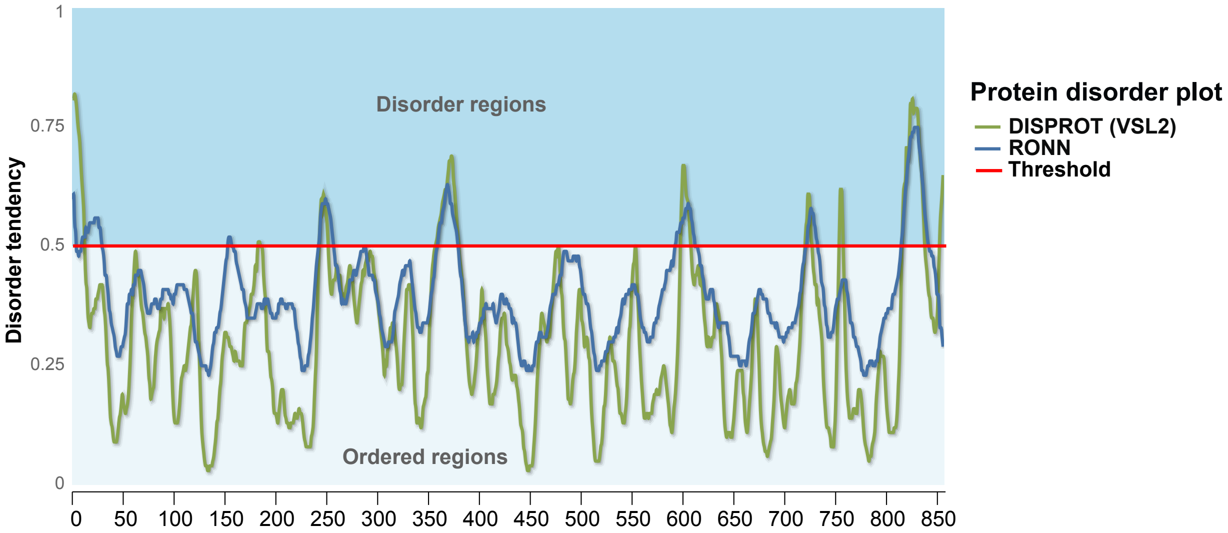

1
0.75
0.5
0.25
0
Disorder tendency
Disorder regions
Ordered regions
0
50
100
150
200
250
300
350
400
450
500
550
600
650
700
750
800
850
Protein disorder plot
DISPROT (VSL2)
RONN
Threshold
